# Robust Removal of Slow Artifactual Dynamics Induced by Deep Brain Stimulation in Local Field Potential Recordings using SVD-based Adaptive Filtering

**DOI:** 10.1101/2023.04.07.536086

**Authors:** Nooshin Bahador, Josh Saha, Mohammad R. Rezaei, Utpal Saha, Ayda Ghahremani, Robert Chen, Milad Lankarany

## Abstract

Deep brain stimulation (DBS) is widely used as a treatment option for patients with movement disorders. In addition to its clinical impact, DBS has been utilized in the field of cognitive neuroscience wherein the answers to several fundamental questions underpinning the mechanisms of neuromodulation in decision making rely on how a burst of DBS pulses, usually delivered at clinical frequency, i.e., 130 Hz, perturb participants’ choices. It was observed that neural activities recorded during DBS were contaminated with stereotype large artifacts, which lasts for a few milliseconds, as well as a low-frequency (slow) signal (∼1-2 Hz) that can persist for hundreds of milliseconds. While the focus of the most of methods for removing DBS artifact was on the former, the artifact removal of the slow signal has not been addressed. In this work, we propose a new method based on combining singular value decomposition (SVD) and normalized adaptive filtering to remove both large (fast) and slow artifacts in local field potentials recorded during a cognitive task in which bursts of DBS were utilized. Using synthetic data, we show that our proposed algorithm outperforms four commonly used techniques in the literature, namely, (1) Normalized least mean square adaptive filtering, (2) Optimal FIR Wiener filtering, (3) Gaussian model matching, and (4) Moving average. The algorithm’s capabilities are further demonstrated by its ability to effectively remove DBS artifacts in local field potentials recorded from the subthalamic nucleus during a verbal Stroop task, highlighting its utility in real-world applications.

## 1. Introduction

The impact of deep brain stimulation (DBS) on local field potential (LFP) recordings can vary depending on the pattern of pulses used. A recent research revealed that multi-pulse stimulation-induced artifacts can persist for up to 1.5 milliseconds [1]. However, earlier studies have found that DBS artifacts can contaminate recorded signals for much longer periods, potentially lasting tens or hundreds of milliseconds [2]. Additionally, another work noted that the shape of DBS artifacts can be influenced by factors such as the shape and impedance of electrodes, the distance between the stimulation and recording sites, as well as the amplitude, frequency, and duration of stimulus pulses [3].

Several filtering techniques have been proposed for removing artifacts from LFP recordings in the literature. These include a low-pass filter [4], a notch filter [5], and band-pass filters [6]. Specifically, [4] focused on removing high-frequency artifacts generated by DBS (130 Hz) by implementing a low-pass filter with a corner frequency of 50 Hz using Butterworth coefficients. Another study [5] reported that no biomarkers for Parkinson’s disease were found in the frequency band between 125 and 155 Hz and thus a high-order (8th) Chebyshev notch filter with a center frequency of 140 Hz and stopband between 125 and 155 Hz was used to suppress artifacts during DBS. A band-pass filter with a frequency band between 100 Hz and the Nyquist frequency of 211 Hz was used to eliminate all stimulation interference [6].

Template subtraction is a commonly used method for eliminating DBS artifacts from LFP recordings. Several studies have explored different methods of implementing this technique [7]. They used thresholding to identify DBS events and then located segments of the LFP that may have been affected by artifacts by analyzing the minimum and zero-crossing points surrounding each detected event. They then employed an ensemble empirical mode decomposition (EEMD) algorithm to decompose the segment into intrinsic mode functions (IMF) and reconstructed the DBS template through a weighted summation of these IMFs, with greater weight given to IMFs with higher frequencies. Another work [8] took a different approach, first detrending the LFP signal by eliminating the residue of the IMF set, then segmenting DBS pulses based on peak points, creating an average artifact shape derived from all pulses, which was then smoothed using a moving average filter to reconstruct the artifact template. Similarly averaging all the stimulus artifact segments and a short time period of 0.4ms around them was used to construct the template of the stimulus artifact [9]. Although template matching is an effective method to identify specific patterns of interest, the variability of DBS artifacts may introduce additional noise into the data, making it difficult to accurately match templates and identify the desired signal.

A technique for removing DBS artifacts from LFP recordings known as detrending uses empirical mode decomposition (EMD) [10]. The approach involved identifying the last intrinsic mode function (IMF) as a trend and removing it to generate a detrended signal. DBS events were then considered as a waveform composed of a set of sine waves, and the amplitude, frequency, and phase of these waves were initialized by visually inspecting the power spectrum. These parameters were then iteratively removed from each segment of the contaminated recording. Another method of removing DBS artifacts is dynamic averaging of small groups of consecutive segments [11, 12]. Segments that are temporally close to each other have similar DBS artifact shapes, and therefore, artifacts can be removed by taking the average of small groups of consecutive segments [11, 12]. Similarly, DBS pulses was detected by amplitude thresholding and took an average of neighboring segments before and after each pulse to generate the stimulus artifact template, which was then subtracted from the raw data at the location of each pulse [13, 14].

Another method for filtering DBS artifacts from LFP recordings is based on temporal decomposition using mutual information between independent components and a reference signal of the DBS [15, 16]. The signals were decomposed into temporally independent components using independent component analysis (ICA) and then computed the mutual information (MI) between the DBS artifact and each of these individual components to determine how much information is shared between them. Finally, components that showed high mutual information were removed. Another technique for removing DBS artifacts is space separation to extract artifact subspace[17-19]. This approach assumes that the DBS component and the original signal are approximately orthogonal to each other, thus, the raw signal is projected into the orthogonal components using signal space projection (SSP), and the DBS artifact is identified and projected out. Eigenvalue decomposition (EVD) of the covariance matrix computed from concatenated epochs including DBS events was performed, and then projected onto orthogonal signal and artifact subspaces. The segments including DBS artifacts were then projected out of the noise subspace. Another study [21] also extracted eigenvectors that span the subspace of DBS artifact at a specific frequency.

Principle-component analysis (PCA) was also employed to identify orthogonal bases that effectively captured the majority of the variance in the data, and to eliminate eigenspaces associated with artifacts [22]. They demonstrated that this approach was more effective at removing deep brain stimulation (DBS) artifacts than alternative techniques such as signal space separation. A different method was employed using thresholding to detect DBS artifact peaks, and subsequently setting samples within the affected period to zero [23]. The missing data was then interpolated by utilizing the neighboring samples that preceded and followed the contaminated period. Other solutions proposed to address the issue of DBS-artifact in LFP recordings include: Adaptive filtering [24], Linear Wiener filtering[25], template subtraction using an adaptive shape based on the Euclidean median of k-nearest neighbors[26], Weighted moving average template subtraction, in which the template is estimated as a weighted average of a limited number of neighboring pulses[27], A template-based subtraction method that utilizes past artifact samples and linear regression[28], Interpolation techniques, including Linear proposed [29], Gaussian [30], and Cubic Spline [31], A symbiotic combination of front-end and back-end template subtraction[32], Polynomial subtraction of power spectral density in the frequency domain to mitigate low-frequency distortions in LFP arising from impedance mismatch during DBS therapy[33]. Despite the effectiveness of these techniques in filtering unwanted noise, there is always a risk of over-filtering, which means that these methods may remove not only unwanted noise but also important information from the signal. This can be particularly problematic for slow wave activities in LFP signals, which can contain important information about brain activity and function.

As previously mentioned, methods such as template matching, may introduce additional noise to the data due to the varying nature of DBS artifacts. While techniques such as adaptive filtering, FIR Wiener filtering, and moving average techniques can effectively preserve fast oscillations, there is a risk of over-filtering, which may result in the removal of important information from the critical slow wave activities in LFP signals. In the present study, we develop a hybrid filtering method that utilizes the strengths of multiple techniques, including SVD and adaptive filtering, to eliminate both sharp spikes and slow wave artifacts.

## 2. Method

The proposed SVD-based Adaptive Filtering technique for removing DBS-induced slow and fast dynamics is conceptually illustrated in Figure 1. These dynamics can greatly impact the processing of LFP and EEG signals during cognitive tasks. In this study, data recorded from patients with Parkinson’s disease (PD) performing a verbal Stroop task was used. As demonstrated in Figure 1, DBS modulates the neural activities of subthalamic neurons while LFP and EEG signals are recorded from multiple electrodes. Through the application of the SVD-based Adaptive Filtering algorithm, the artifacts observed during and after DBS pulses were effectively removed. (See Section 2.2.1 for further information on the experimental data).

**Fig.1.**
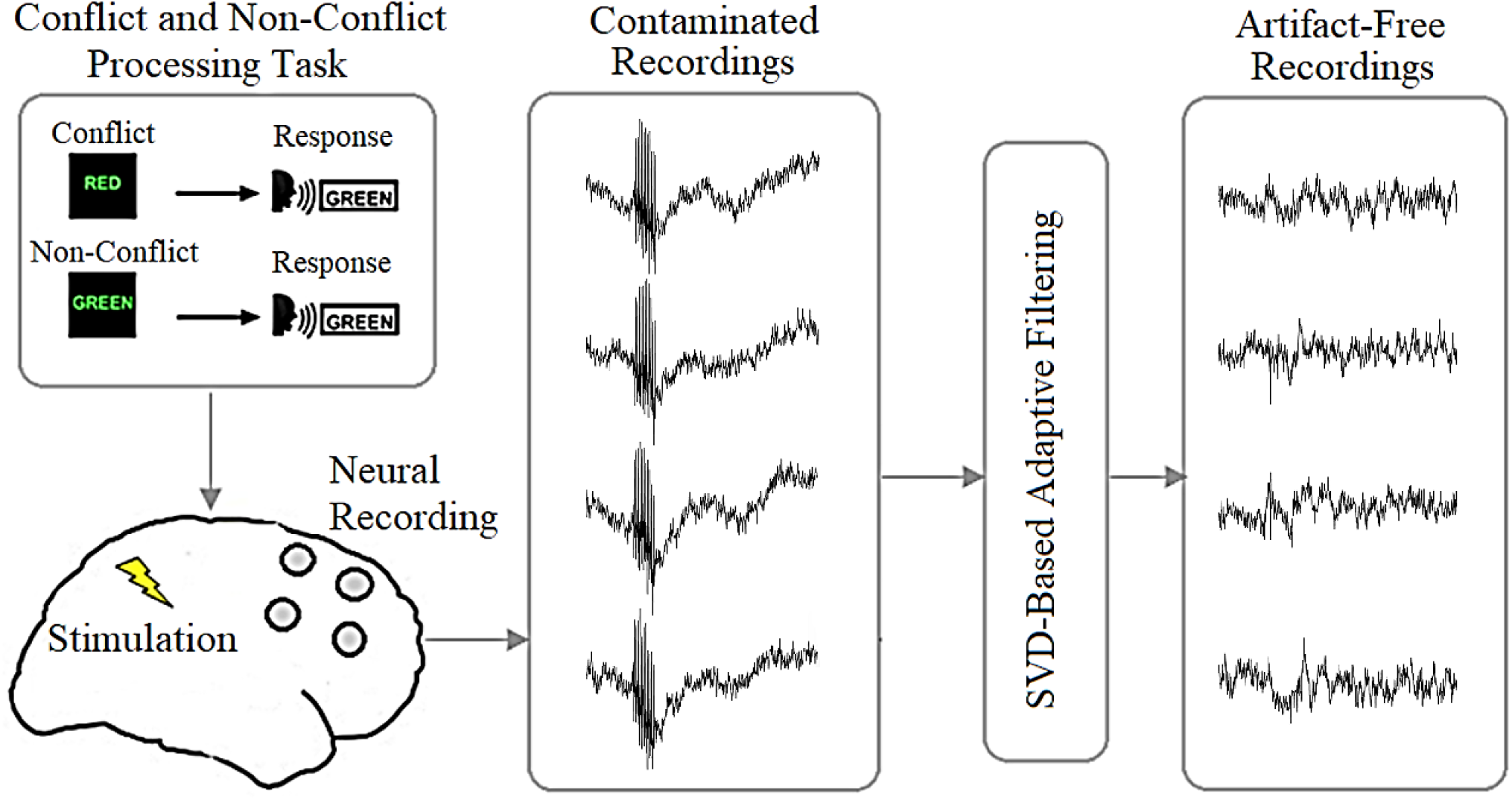
A Conceptual Framework for SVD-based Adaptive Filtering of DBS-Induced Artifacts during a Verbal Stroop Task: This figure illustrates the conceptual framework of the SVD-based adaptive filtering algorithm for removing both high- and low-frequency artifacts induced by Deep Brain Stimulation (DBS) during a verbal Stroop task. The final window in the figure presents the outcome of the algorithm, where we can see the signal without DBS artifact, and therefore yielding a more accurate measurement of neural activity related to cognitive processes.

### 2.1. Data

The present study includes an examination of both synthetic and experimental data sets.

#### 2.1.1. Synthetic Data

The synthetic dataset comprised of a sine wave, colored noise, and a DBS artifact template (as illustrated in Figure 2). A 1 Hz sine wave with the same amplitude as the original signal was included to simulate the slow-wave artifact that can be induced by movement. Additionally, a Gaussian white noise was generated and passed through an IIR digital filter of order 10 to produce brown noise.

**Fig.2.**
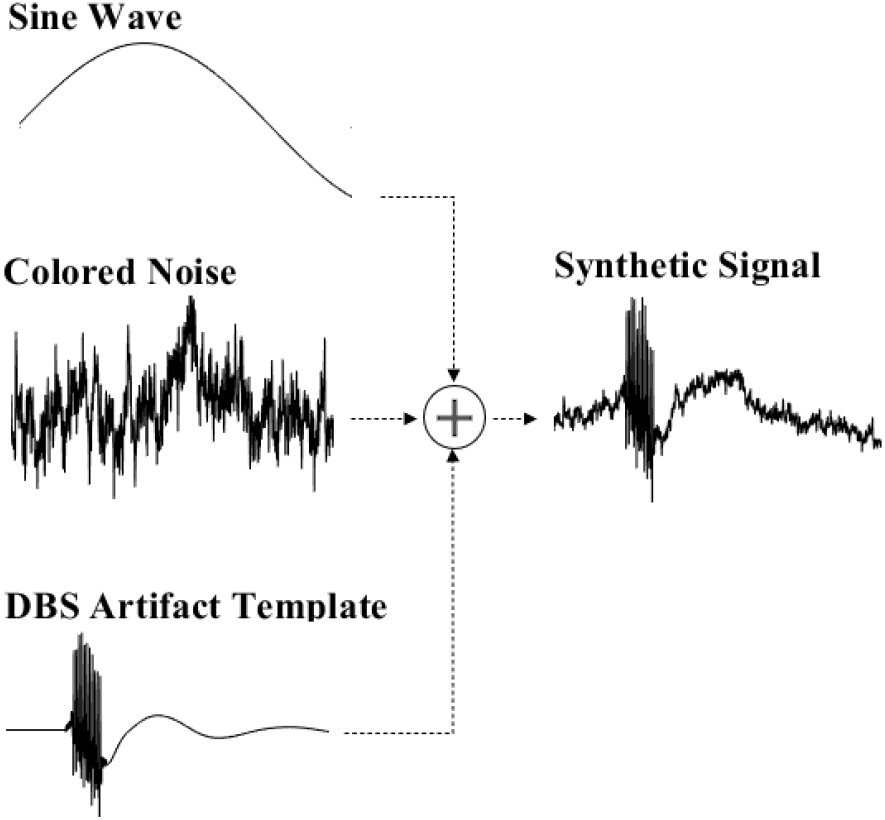
Synthetic signal generation process: This figure illustrates the procedure for creating synthetic signals, which consist of a combination of a sine wave representing slow-wave movement artifacts, colored noise representing neural activity, and DBS artifacts modeled on a stimulus artifact template.

#### 2.1.1. Experimental Data

For the experimental portion of the study, data was collected from patients with Parkinson’s disease who were receiving trains of DBS during specific periods of a Stroop task. The task involved displaying a color word on a screen and asking the patients to name the ink color of the word into a microphone. In the conflict trials, the word and ink color did not match, while in the non-conflict trials, they did match [34].

Four different stimulation conditions were used, with each condition applied randomly to one trial out of every four. These four conditions were: no stimulation, Ready period, Early response, and Late response. Each patient completed a total of 240 trials, with event-related stimulation applied during each trial after a certain number of test trials had been completed. Bilateral STN LFPs and scalp EEG were recorded during the trials, using a sampling frequency of 5 kHz and a filtering range of 1-1000 Hz. The data was down-sampled to 1000 Hz for analysis. Only trials with response times between 0.3 and 1.8 seconds were included in the analyses [34].

The trains of pulses used for stimulation consisted of 11 pulses with a duration of approximately 80 milliseconds, a frequency of 130 Hertz, a monophasic waveform, and a pulse width of 100 microseconds. The stimulation intensity for event-related deep brain stimulation (DBS) was 1.92 ± 0.76 milliamperes (mA). The trigger pulses for stimulation were generated using Spike2 software from Cambridge Electronic Design in the UK and were delivered through a constant current stimulator from Digitimer in Welwyn Garden City, Hertfordshire, UK [34].

Neurophysiological mapping was employed to confirm the stimulated contact’s location. Intraoperative recordings were used to identify the ventral border of the STN by detecting a change in the pattern of cell firing, which was then used to determine the final target for the DBS electrode implantation. The STN’s top and bottom margins were determined with reference to the DBS electrode’s final position (DBS 3387 electrode, Medtronic, Minneapolis), as reported in the operative notes. The location of each contact relative to the STN was estimated using the electrode’s known dimensions, i.e., four 1.5mm contacts spaced 1.5mm apart [34].

### 2.2. Benchmark DBS-artifact suppression techniques in this study

For the comparison of the performance of the proposed technique, four existing methods were used as benchmark techniques. These methods were normalized least mean square adaptive filtering [35], optimal FIR Wiener filtering [36], Gaussian model matching [37], and moving average techniques [27].

#### 2.2.1. Normalized least mean square (NLMS) adaptive filter algorithm

The main structure of a NLMS filter consists of a FIR filter with coefficient update procedure in which the coefficients are updated so that the difference between the output and the reference signal becomes minimum.

The NLMS filter is defined by the following equations.

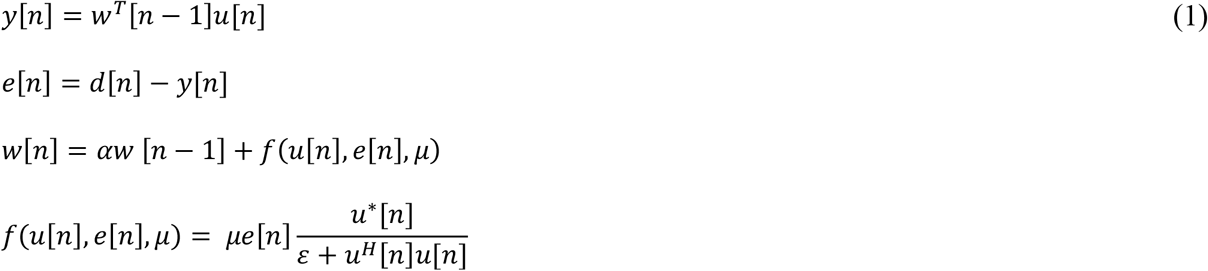

where n is the current index of temporal sample. u is input signal. *u*^*^ is complex conjugate of input signal. w is filter weight. y is output signal. e is estimation error. d is the reference signal. *μ* is adaptation step size. *ε* is stability constant [38].

#### 2.2.2. Optimal FIR Wiener filter

In the context of artifact subtraction using Optimal FIR Wiener filter, the target signal contaminated by artifact, and the estimate of artifact provided by the Wiener filter using reference signal. The optimal filter satisfies the Wiener-Hopf equation given by following equation [39].

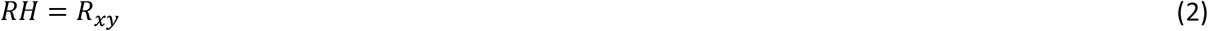

where R, H and *R*_*xy*_ are respectively the symmetric Toeplitz autocorrelation matrix of the input signal, impulse response of the optimal filter, and cross-correlation between the input and the desired output. The matrix of R is of the form given by:

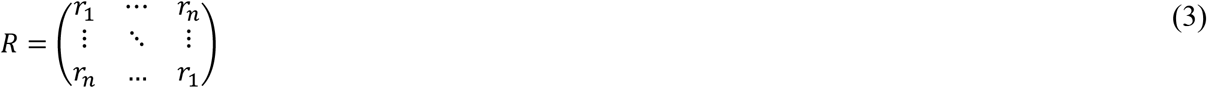

#### 2.2.3. Gaussian model matching

The global peaks in the signal are fitted using Gaussian model which is given by

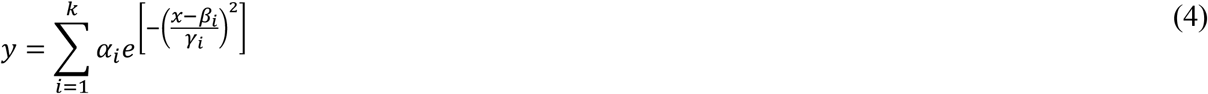

where *α, β, γ* and k are respectively the amplitude, centroid, peak width and the number of peaks to fit.

This fitted model is then used as the artifact template and subtracted from original signal to provide a clean version of signal.

#### 2.2.4. Moving average

The artifact template is generated based on the local k-sample average values, where each average is calculated over a sliding window of length k across neighboring samples of original signal.

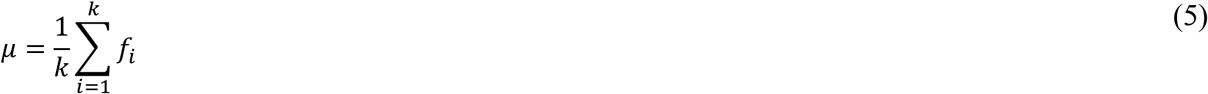

The filtered signal is then obtained by template subtraction from the original signal.

### 2.3. Proposed SVD-based adaptive filtering

The SVD Adaptive Filtering algorithm is a method for removing slow component of DBS artifact from an input signal while also reducing the presence of sharp DBS peaks. This filtering includes two separate modules for removing slow and fast waves of DBS artifact (DBS artifact starts with fast and sharp transients followed by a slow wave).

The process begins by initializing the input signal. The algorithm then generates a set of circularly shifted versions of the input signal by repeatedly shifting the elements a certain number of positions. These shifted signals are stored and then combined to create a matrix. Next, the algorithm performs SVD to obtain the matrices U, S, and V. The diagonal matrix S is extracted, and any singular values that are more than 90% of the maximum singular value are set to zero (Figure 3). Using the modified S matrix, along with U and V, the algorithm reconstructs the signal. A reference signal is then calculated by finding the moving average of the corresponding values, with a window size of 20. Starting from the DBS time until the end of the array, the values of reconstructed signal are replaced with the reference signal. An adaptive filtering algorithm using the Least Mean Squares (LMS) method is then applied to extract the high frequency component and suppress the DBS peaks (Figure 4). This process is repeated twice more to further reduce the amplitude of these peaks.

**Fig.3.**
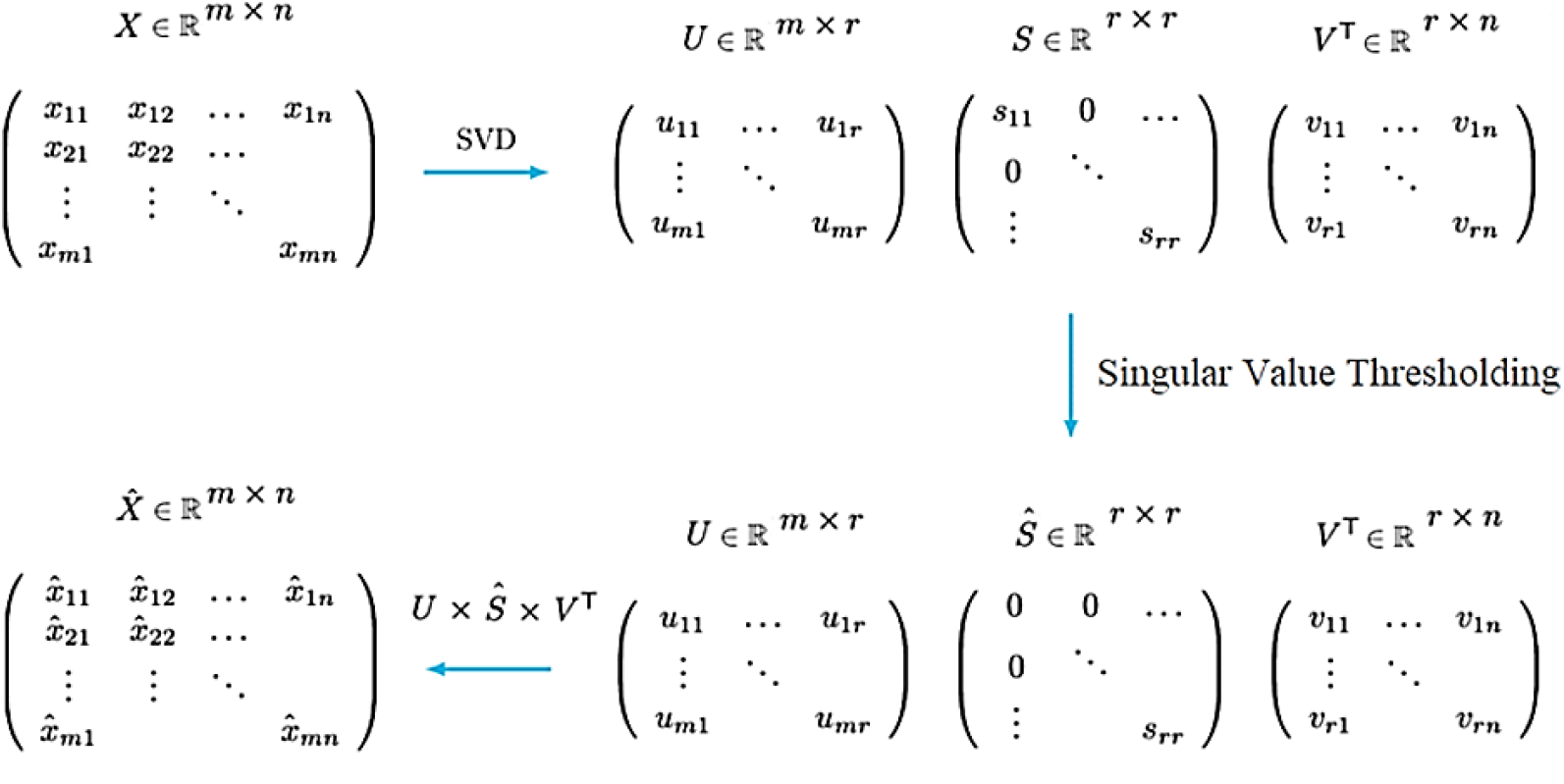
Signal reconstruction based on singular value decomposition

**Fig.4.**
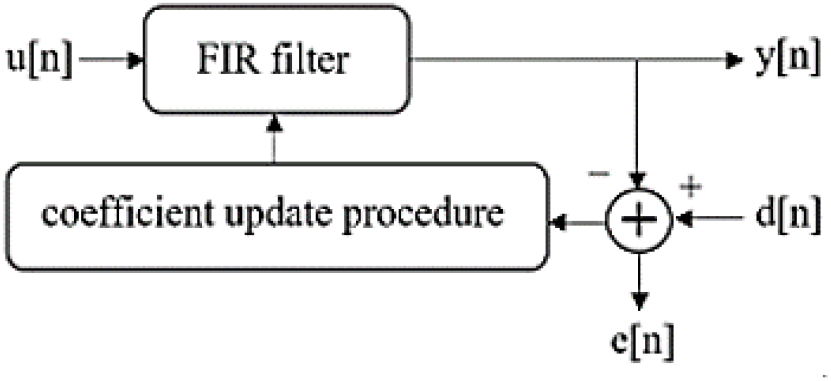
Adaptive filtering procedure

Here’s the algorithm that describes the above steps in the proposed technique:

#### Algorithm

SVD Adaptive Filtering

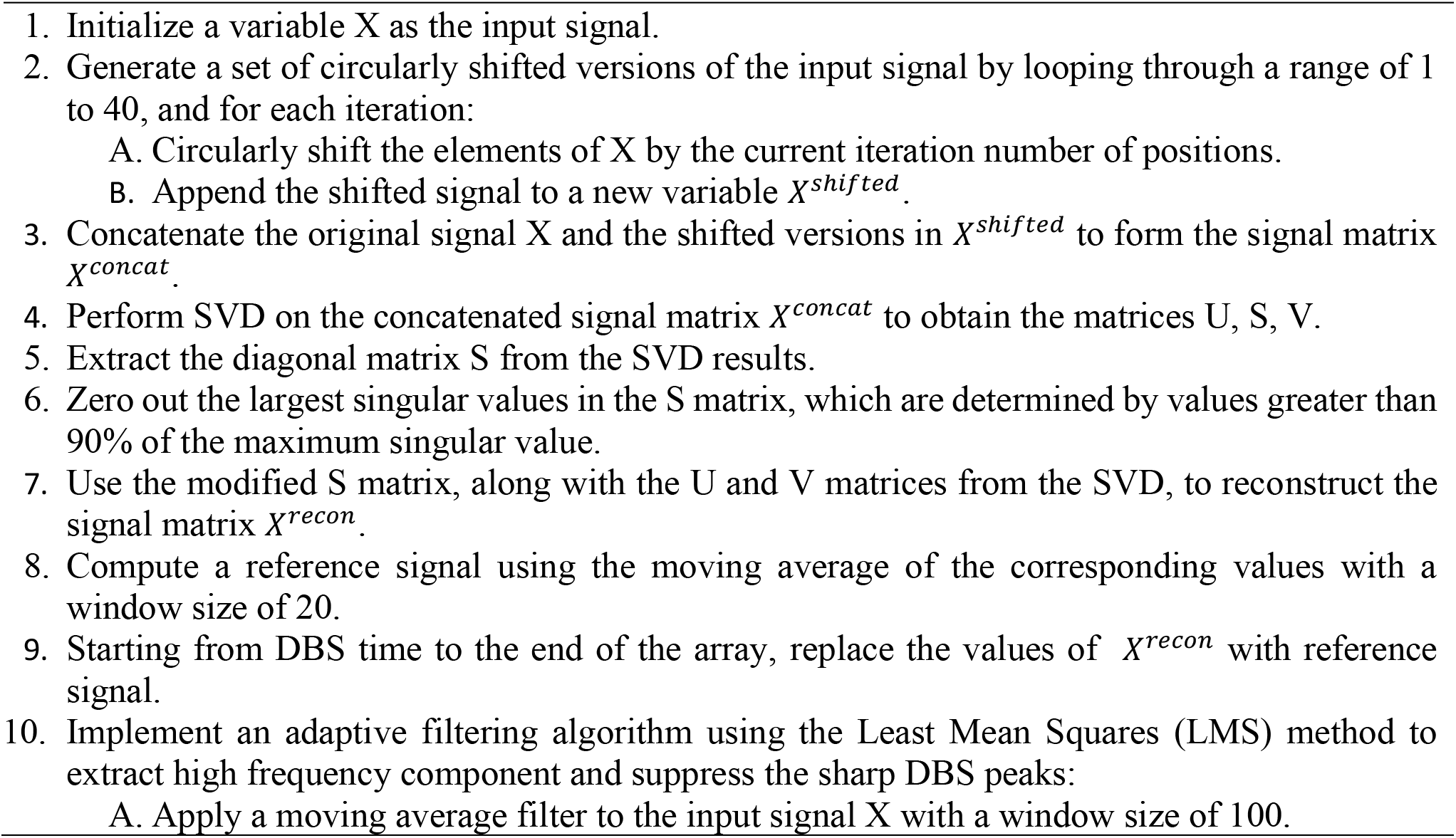
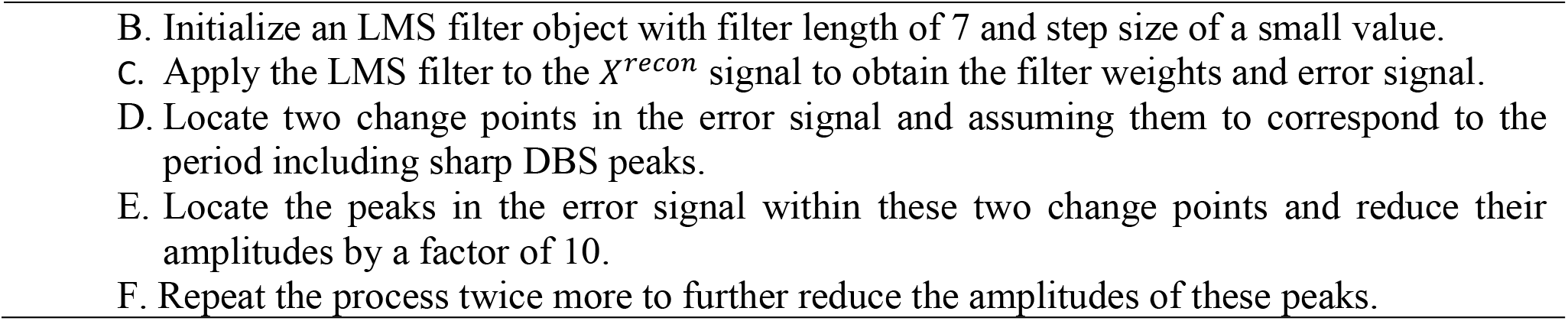

## 3. Results

### 3.1. Validation of artifact removal algorithms on synthetic LFP signal

Figure 5 presents a comparison of the contaminated signals for positive and negative pulses. The first sample (left) illustrates the contaminated signal when the pulse is negative, while the second sample (right) shows the contaminated signal when the pulse is positive. This comparison highlights the effect of DBS pulse polarity on the contamination of the signal.

**Figure 5.**
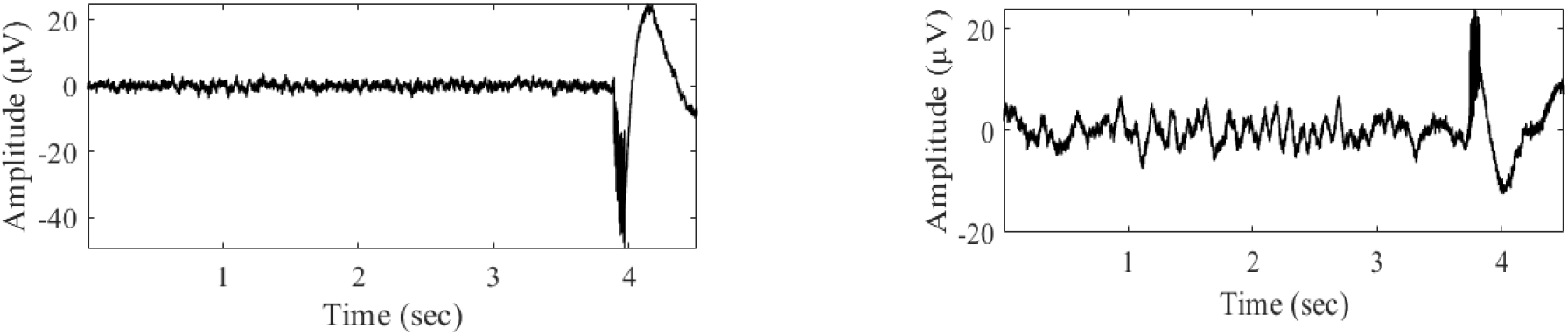
The effect of reversing the polarity of the stimulus on the slow wave component of DBS artifacts: When the stimulus is reversed, the artifact seen in the is reversed or mitigated.

The time-frequency analysis in Figure 6 was conducted to assess the preservation of the original signal’s temporal and spectral information after DBS artifact removal. The availability of the clean version of synthetic signals made it easy to evaluate the performance of the artifact removal algorithm. By visually comparing the time-domain signals before and after the removal of DBS artifacts, it is evident that the algorithm not only effectively eliminated the sharp spikes caused by DBS, but also the slow activities induced by both the DBS artifact and sine wave. Additionally, by observing the small fluctuations in amplitude of the original signal, it can be seen that the filtered version closely resembles the original signal. The comparison of the spectrogram plots in Figure 6 between the original and artifact-free signals confirms the preservation of temporal-spectral patterns within the original signal and demonstrates the good performance of the proposed filtering technique.

**Figure 6.**
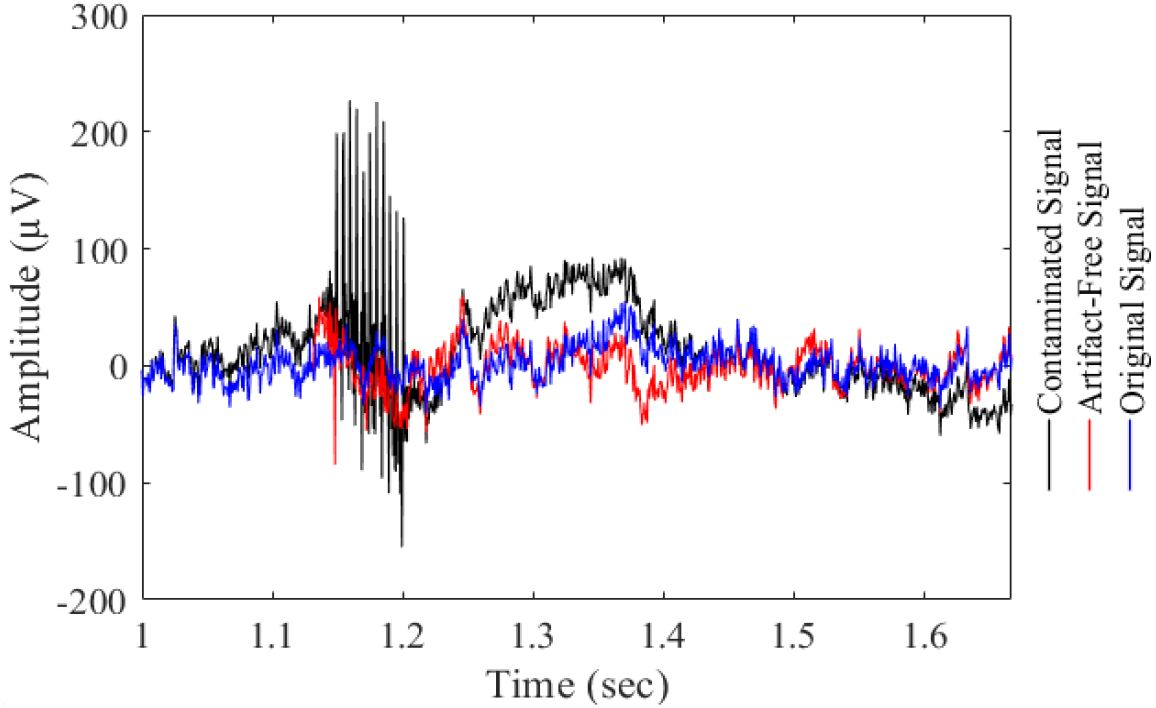
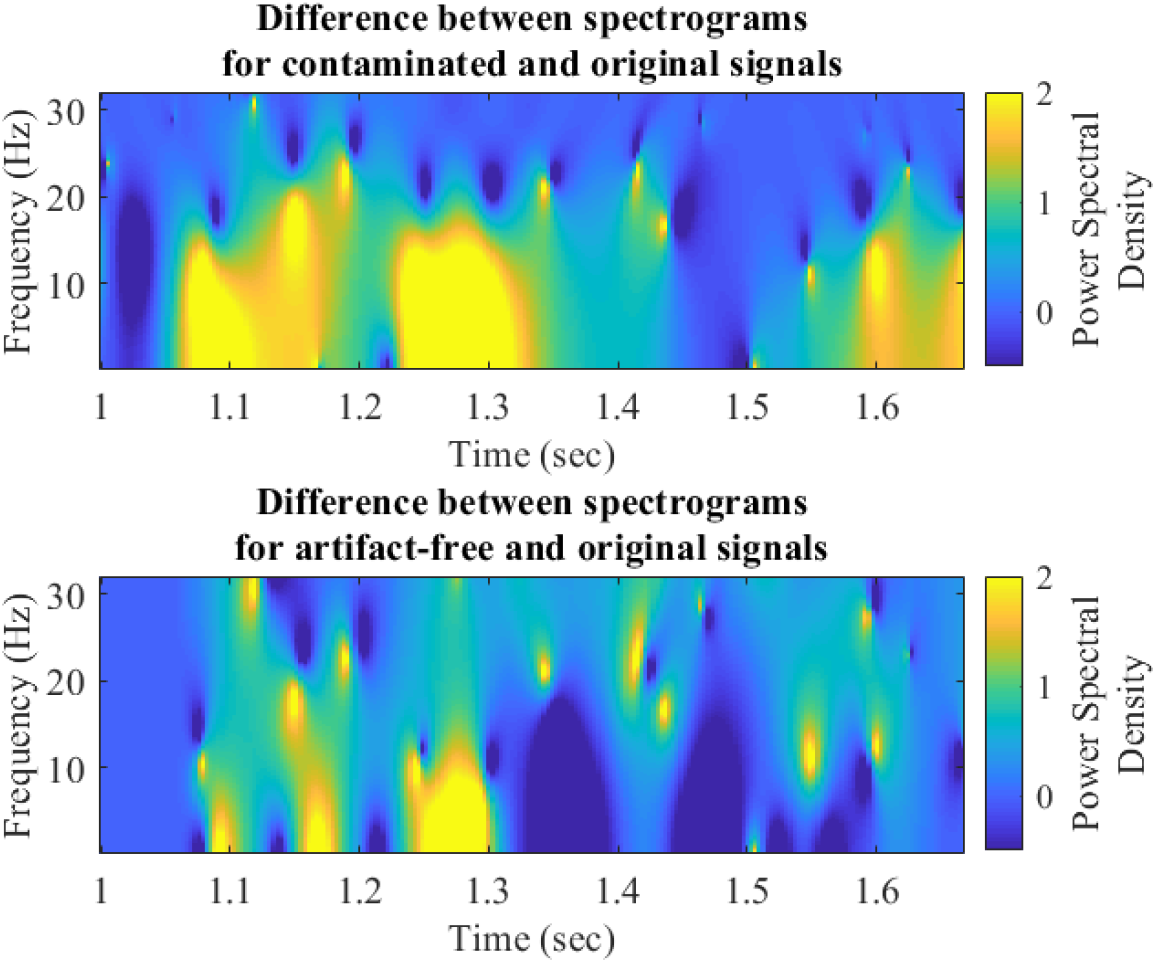
The first subplot compares the clean synthetic signal after artifact removal (in red) to the original (in blue) and contaminated (in black) synthetic signals. The temporal analysis highlights the effectiveness of the artifact removal process in restoring the integrity of the clean signal. The second and third subplots show the difference in spectrograms between the original and contaminated signals and the original and artifact-free signals, respectively. The spectral analysis illustrates the reduction of artifacts in the time-frequency domain as a result of the artifact removal process.

Mean-squared error (MSE) between the original synthetic signal and contaminated one was compared to MSE between the original synthetic signal and artifact-free one in time domain (Table 1). The percentage difference obtained by SVD filtering were 95.26% which outperformed other techniques. Adaptive filtering came in second place by percentage difference of 46.99%. Based on these results, it was hypothesized that combining SVD with adaptive filtering may improve the suppressing of both DBS and motion artifacts. This hypothesis was tested on real data and the results was presented in following subsection.

**Table 1.** Time domain comparison of MSE values

| Technique | Mean-squared error (MSE) between the<br>original signal and |  | Percentage<br>Difference<br>in MSE % |
| --- | --- | --- | --- |
|  | Contaminated Signal | Artifact-Free Signal |  |
| SVD | 1,230 | 58.57 | -95.26 |
| Gaussian model matching | 1,230 | 1,540 | +24.91 |
| FIR wiener filtering | 1,230 | 671.55 | -45.76 |
| Adaptive filtering | 1,230 | 656.27 | -46.99 |
| Moving average | 1,230 | 1,400 | +16.01 |

### 3.2. Validation of artifact removal algorithms on experimental LFP signal

The Figure 7 displays the evaluation of various filtering techniques, including SVD (Figure 7 A), adaptive filtering (Figure 7 B), FIR Wiener filtering (Figure 7 C), Gaussian model matching (Figure 7 D), and moving average (Figure 7 E) for suppressing artifacts in DBS signals, with the aim of identifying the most effective combination of techniques for eliminating sharp spikes and slow waves caused by DBS pulses. Figure 7E shows that the use of a moving average window smoothed out most slow activities, including non-artifactual components, and yet sharp spikes induced by DBS remained present in the filtered signal. Similarly, adaptive filtering in Figure 7B and FIR wiener filtering in Figure 7C flattened out artifactual and non-artifactual slow activities without removing sharp spikes. In contrast, the use of a gaussian model matching filter in Figure 7D introduced unwanted noise into the signal. It is clear from these results that the use of a single technique, such as moving average filtering, adaptive filtering, or FIR wiener filtering, is not sufficient to effectively remove both fast and slow artifactual components without affecting the useful neural information. Instead, the most effective approach would be to utilize the strengths of multiple techniques, such as the SVD technique (Figure 7 (A)) for reproducing clean slow wave activities in the original signal and adaptive filtering for reproducing fast oscillations. In this way, both sharp spikes and slow wave artifacts induced by DBS can be effectively eliminated.

**Figure 7.**
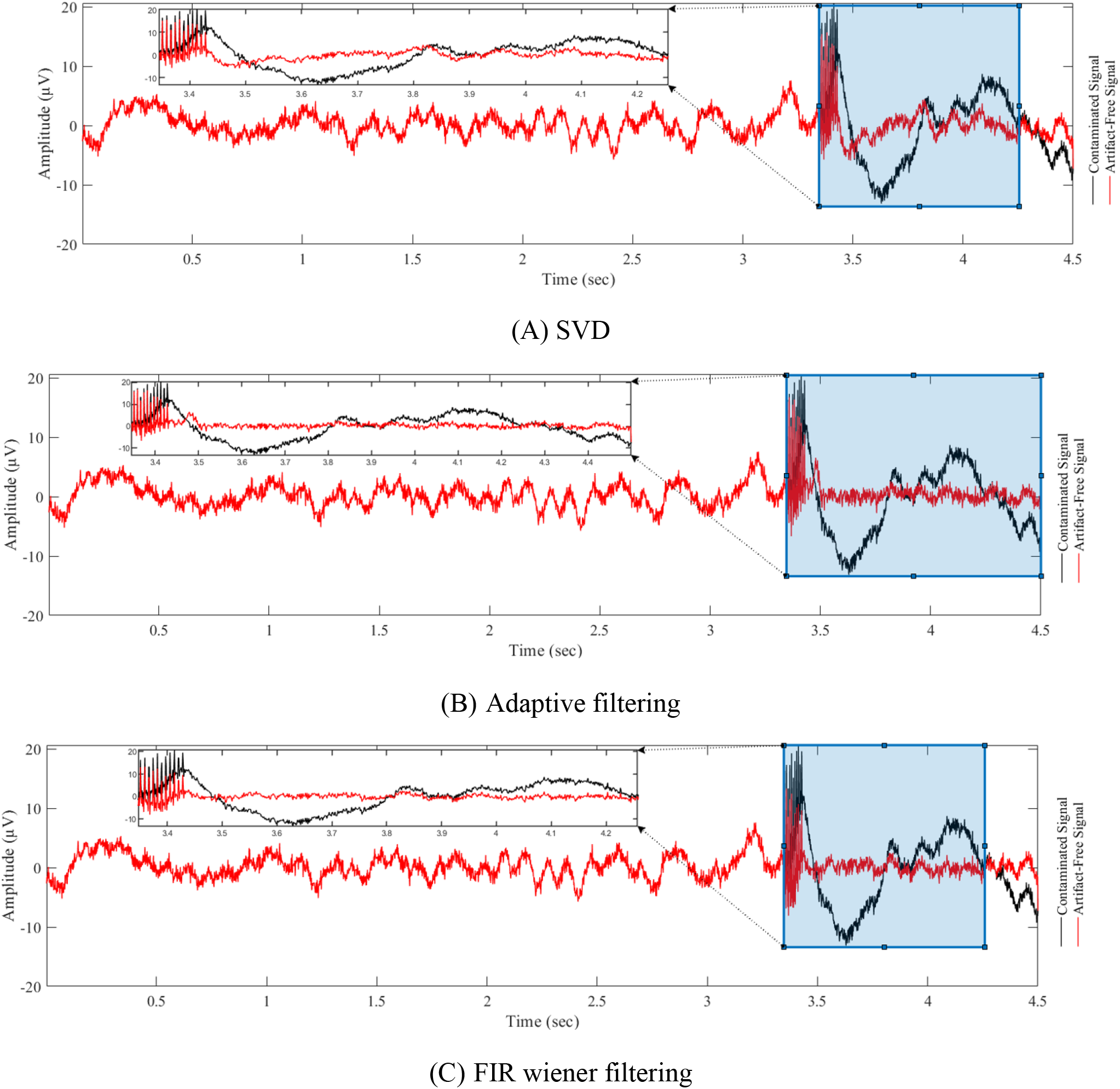
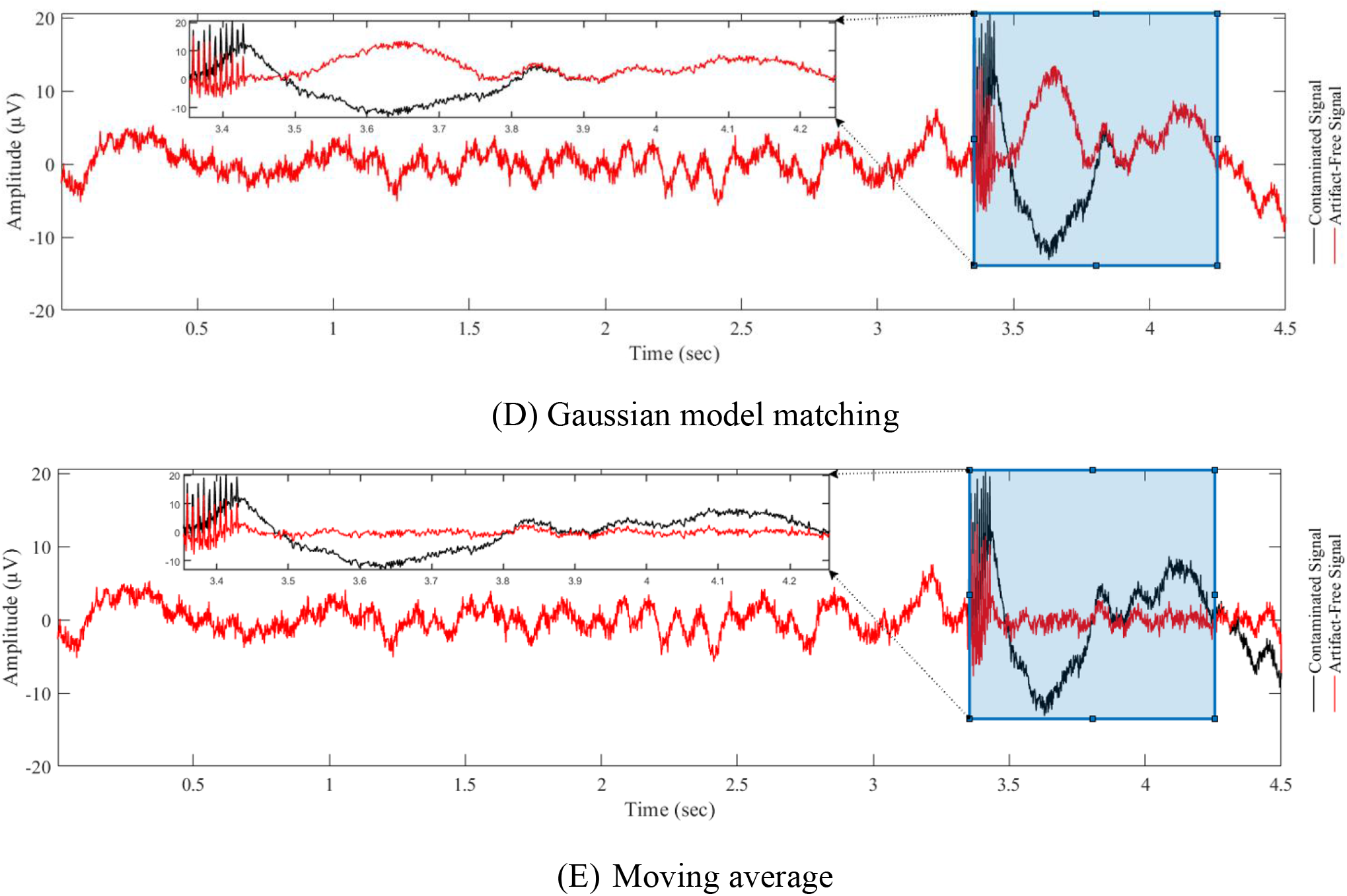
Evaluation of various techniques for suppressing artifacts in DBS signals. The aim of this comparison was to identify the most effective combination of techniques for eliminating both sharp spikes and slow waves caused by DBS pulses. A comprehensive analysis of different filtering methods were considered to provide insight into ways to improve the overall quality of DBS filtering.

Figure 8 presents the results of an SVD-based adaptive filtering technique applied to remove DBS artifacts. The subplot A of the figure demonstrates the effectiveness of the technique in preserving neural activity while effectively filtering out the sharp spikes and slow bumps associated with DBS. The low and high frequency components induced by neural activity remain untouched, while the DBS artifacts are successfully removed. The subplots of B and C provide additional support for the technique’s effectiveness through the use of spectrograms. A spectrogram is a visual representation of the frequency content of a signal over time, and it provides a powerful tool for analyzing and interpreting signals, particularly those that vary over time. The spectrograms in these subplots demonstrate the effectiveness of the SVD-based adaptive filtering technique in removing the slow wave activities induced by DBS. By eliminating these artifacts, the spectrograms reveal a clean and accurate representation of neural activity. Specifically, the spectral content of the filtered signal is seen to closely match the spectral content of the original neural activity, with a significant reduction in the spectral content of the DBS artifacts. This is a strong indication that the SVD-based adaptive filtering technique is effectively removing these artifacts, while preserving the underlying neural activity.

**Figure 8.**
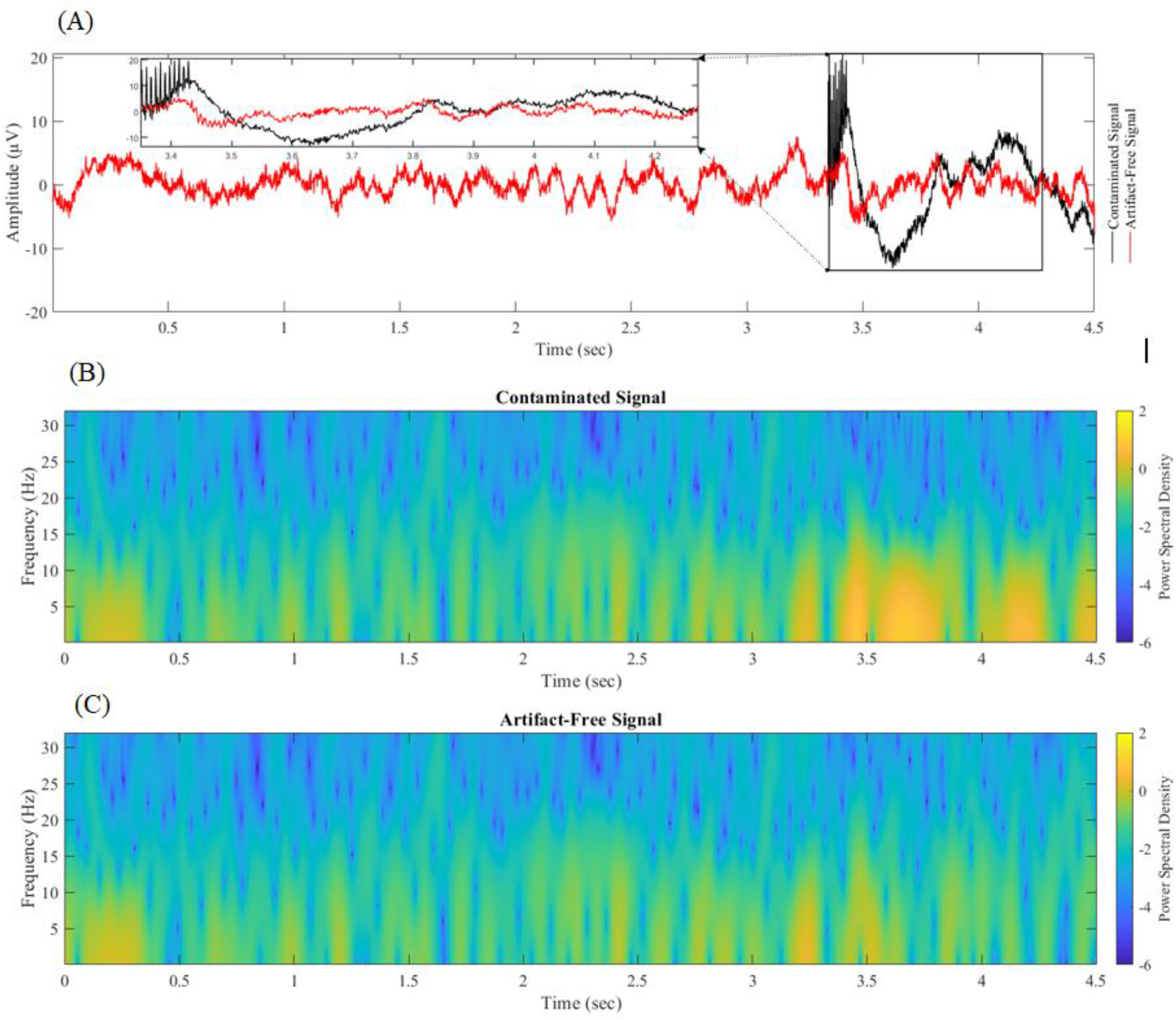
SVD-based Adaptive Filtering for DBS Artifact Removal. This figure illustrates the effectiveness of combining SVD with adaptive filtering for removing deep brain stimulation (DBS) artifacts. The subplot A compares the contaminated signal with the filtered version in the time domain, with the enlarged window highlighting the beginning of DBS. The subplots B and C provide a spectral analysis of the contaminated and filtered signals, respectively, through the use of spectrograms. The SVD-based adaptive filtering technique is able to effectively remove DBS artifacts while preserving the underlying neural activity, as demonstrated by the clean version of the signal and the consistent spectral content in the filtered version of the signal.

## 4. Conclusion

Neurostimulation has emerged as a cutting-edge technology for the treatment of neurological disorders, offering great potential for closed-loop neuromodulation. However, recorded neural signals such as LFPs are often contaminated by artifacts induced by stimulation pulses. We observed that a short DBS burst induce both fast (sharp, lasting for a few msec) and slow (lasting for hundreds of msec) artifacts.

Here we proposed a solution using SVD-based adaptive filtering. To address the issue of overfiltering with adaptive filtering, an additional module was included in the study to separate meaningful slow wave activities from artifacts. This module reproduced the slow wave activities in LFP recordings by applying the SVD and considering singular values less than a specific threshold. Our synthetic results showed that the proposed technique exhibits superior performance compared to existing methods, successfully recovered both high and low-frequency neural information without increasing noise levels. Using LFPs recorded from STN during verbal Stroop task, we demonstrated that SVD-adaptive filtering successfully removed slow artifactual dynamics induced by a short burst of DBS.

The proposed adaptive filtering approach improved detection of neural oscillations in a short time period after DBS pulses, which in turn can be useful for enhancing efficacy of closed-loop neuromodulation devices. By effectively suppressing DBS artifacts from LFP recordings, we anticipate that the proposed SVD-based adaptive filtering method will provide a valuable tool for researchers and practitioners in the field of neurostimulation. In future work, we will improve the implementation of the algorithm for real-time applications.

